# Toward the pathogenicity of the *SLC26A4* p.C565Y variant using a genetically driven mouse model

**DOI:** 10.1101/2020.04.29.067918

**Authors:** Chin-Ju Hu, Ying-Chang Lu, Ting-Hua Yang, Yen-Hui Chan, Cheng-Yu Tsai, I-Shing Yu, Shu-Wha Lin, Tien-Chen Liu, Yen-Fu Cheng, Chen-Chi Wu, Chuan-Jen Hsu

## Abstract

Recessive variants of the *SLC26A4* gene are a common cause of hearing impairment worldwide. In the past, cell lines and transgenic mice have been widely used to investigate the pathogenicity associated with the *SLC26A4* variants. However, discrepancies in the pathogenicity between humans and cell lines or transgenic mice have been documented for some of the *SLC26A4* variants. For instance, the p.C565Y variant, which has been reported to be pathogenic in humans, did not exhibit functional pathogenic consequences in cell lines. To address the pathogenicity of p.C565Y, we used a genotype-based approach in which we generated knock-in mice heterozygous (*Slc26a4*^+*/C565Y*^), homozygous (*Slc26a4*^*C565Y/C565Y*^), and compound heterozygous (*Slc26a4*^*919-2A*>*G/C565Y*^) for this variant. Subsequent phenotypic characterization revealed that mice segregating these genotypes demonstrated normal auditory and vestibular functions and normal inner ear morphology and expression of pendrin. These findings indicate that the p.C565Y variant is non-pathogenic for mice and that a single p.C565Y allele is sufficient to maintain normal inner ear physiology in mice. Our results highlight the differences in the pathogenicity associated with certain *SLC26A4* variants between transgenic mice and humans, which should be taken into consideration while interpreting the results of animal studies for *SLC26A4*-related deafness.

## Introduction

Recessive variants of the *SLC26A4* (Gene ID: 5172) gene are considered as a common cause of hereditary hearing impairment (HHI) worldwide [1]. In certain populations, pathogenic *SLC26A4* variants can be identified in approximately 15-20 % patients with HHI [2]. *SLC26A4* encodes for pendrin, a chloride bicarbonate transporter, which is mainly expressed in the thyroid, inner ears, kidneys, lungs, liver, and heart [3, 4]. Recessive *SLC26A4* variants are causal for the occurrence of Pendred syndrome (PS; MIM #274600) and non-syndromic hearing loss, DFNB4 (MIM #600791). DFNB4 is characterized by isolated sensorineural hearing impairment (SNHI), which is usually associated with a common inner ear malformation known as enlarged vestibular aqueduct (EVA; MIM 603545); whereas patients with PS have goiter in addition to EVA [5]. Clinically patients with *SLC26A4* variants, either with the manifestation of DFNB4 or PS, usually suffer from progressive or fluctuating SNHI [6].

To date, more than 400 *SLC26A4* variants have been reported (http://deafnessvariationdatabase.org/). Corresponding to the extensive variation in the severity of clinical phenotypes, different *SLC26A4* variants are shown to be associated with different pathogenic consequences in cell-line experiments [7, 8]. Some variants, such as p.H723R, p.L236P, and p.T721M, are associated with defective protein expression or trafficking, which leads to the accumulation of pendrin either in the cytoplasm or perinuclear region. Some variants, such as p.K369E and p.S166N, have been shown to be associated with normal protein expression of pendrin in the cell membrane but impaired protein function for chloride bicarbonate (Cl^-^/ HCO_3_ ^-^) exchange [7-9]. Interestingly, an enigmatic *SLC26A4* variant, p.C565Y, which was reported to be pathogenic in humans, did not exhibit any functional pathogenic consequences in the cell line. Therefore, it is extremely crucial to clarify the discrepancy in the pathogenicity between humans and cell lines in order to dissect the pathogenic mechanisms related to *SLC26A4* variants.

In addition to the cell-line models, transgenic mice have also been demonstrated to be a powerful tool for investigating the pathogenic mechanisms of *SLC26A4* variants [10-15]. In this study, we generated a knock-in mouse model harboring the p.C565Y variant of *SLC26A4* and then further investigated the associated audiovestibular phenotype and the inner ear pathology.

## Materials and Methods

### Generation of *Slc26a4*^*C565Y/C565Y*^ knock-in mice

Transgenic mice were generated by using the Transgenic Mouse Models Core (TMMC, Taiwan) and the clustered regularly interspaced short palindromic repeat (CRISPR) technology-associated RNA-guided endonuclease Cas9 to mutate the *Slc26a4* gene and further generate the *Slc26a4*^+*/C565Y*^ mouse strain. Specific guide RNAs (sgRNAs) were developed to target exon 15 of the *Slc26a4* gene in the C57BL/6 mouse strain. The sgRNA and CRISPR/Cas9 RNA were delivered simultaneously into the zygote of C57BL/6 mouse to generate the founders. Two male founder mice were obtained, each possessing the p.C565Y (c.1694G>A) variant in the *Slc26a4* gene. After germline transmission of the targeted mutation allele, we were able to generate the congenic *Slc26a4*^+*/C565Y*^ mouse line by repeated backcrossing with the C57BL/6 inbred strain for 6–10 generations. Thereafter, the homozygous mice for the *Slc26a4* p.C565Y variant (*Slc26a4*^*C565Y/C565Y*^) were obtained by intercrossing heterozygous mice (*Slc26a4*^+*/C565Y*^ x *Slc26a4*^+*/C565Y*^) (Fig. 1A and B).

**Figure 1.**
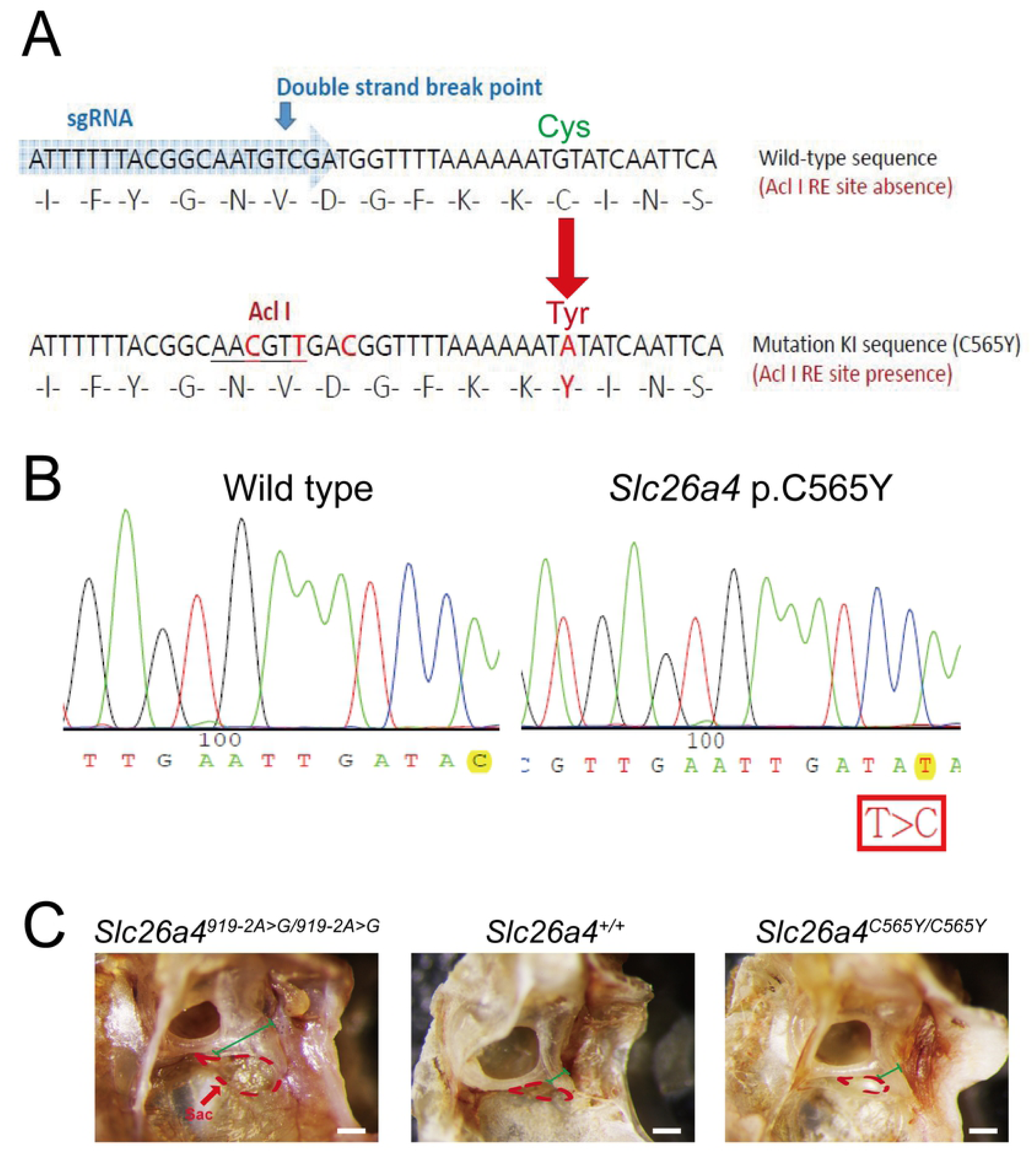
Generation of mice with the *Slc26a4* p.C565Y variant using CRISPR/Cas9. (A) Design diagram. SgRNA for CRISPR/Cas9 and silent mutations for enzyme cutting sites (for checking) were designed to incorporate the mutations into the genome of C57BL/6 mice. To generate the *Slc26a4* p.C565Y variant, the codon “TGT” was mutated to “TAT” (B) Sanger sequencing was performed to confirm nucleotide changes in transgenic mice. The sequence was read in reverse. (C) Gross morphology. The enlargement in the endolymphatic sac (the region delineated by dotted line) and vestibular aqueduct (marked by red line) was observed in the *Slc26a4*^*919-2A*>*G/919-2A*>*G*^ mice and were not observed in *Slc26a4*^+*/*+^ and *Slc26a4*^*C565Y/C565Y*^ mice (bar = 100μm).

Corresponding to the human genotypes, compound heterozygous mice (i.e. *Slc26a4*^*919-2A*>*G/C565Y*^) with both the p.C565Y and c.919-2A>G variants were also generated by intercrossing heterozygous *Slc26a4*^+*/C565Y*^ mice with *Slc26a4*^*919-2A*>*G/919-2A*>*G*^ mice [13]. All the animal experiments were carried out in accordance with the Animal Welfare guidelines and were approved by the Institutional Animal Care and Use Committee (IACUC) of National Taiwan University College of Medicine (approval no. 20160337).

### Auditory evaluations

For audiological evaluations, the mice (postnatal 28 days, P28) were anesthetized and maintained in a head-holder within an acoustically and electrically insulated and grounded test room [13]. We used an evoked potential detection system (Smart EP 3.90; Intelligent Hearing Systems) to measure the thresholds of the auditory brainstem response (ABR) in mice. Click sounds as well as 8, 16, and 32 kHz tone bursts at varying intensity were given to evoke ABRs of subject mice. To determine ABR thresholds, sound pressure levels were used between 15 and 70 dB SPL. The response signals were detected using the subcutaneous needle electrodes.

### Vestibular evaluations

For vestibular evaluations, the mice were subjected to a series of tests (all performed at 8 weeks of age), including the swimming test and the rotarod test. For the swimming test, the swimming performance of the mice was scored from 0 to 3, with 0 being normal swimming and 3 being underwater tumbling [16]. For the rotarod test, the mice were placed on the rotating rod for a maximum of 180 s. The speed of the rods was accelerated from 5 rpm to the maximum speed of 20 rpm for 1 min. The duration for which each mouse was able to sustain on the rotating rod was recorded [17].

### Inner ear histology studies

To perform the light microscopy analysis, tissues were subjected to hematoxylin and eosin (H&E) staining, and the morphology of each sample was examined using a Leica optical microscope (Leica). At first, the inner ear tissues from adult mice (3-month-old) were harvested, and were then fixed using perilymphatic perfusion with 4% paraformaldehyde (PFA) (Bio Basic Inc.,) prepared in phosphate-buffered saline (PBS). The specimens were decalcified for 1 week. Later, the samples were dehydrated and embedded in paraffin. Subsequently, serial sections (7 µm) were obtained and stained with H&E.

For whole-mount studies of mouse inner ear, specimens were prepared as previously described [13]. The tissues were stained with rabbit anti-Myosin-VIIA primary antibody (1:200; Proteus Bioscience Inc.,). Later, the tissue sections were incubated with DAPI (1:5000; Thermo Fisher) and Alexa Fluor 568-conjugated goat anti-rabbit IgG (H+L) secondary antibodies (1:200; Thermo Fisher) at 4°C overnight. After washed with PBS to remove any residual antibodies, the tissue samples were mounted using the ProLong Antifade Kit (Molecular Probes) for 20 min at room temperature. Images of the mice inner ear tissues were captured using a laser scanning confocal microscope (LSM880, Zeiss).

### Expression of pendrin

For pendrin expression analysis, tissue sections were harvested from the inner ears of *Slc26a4*^*919-2A*>*G/C565Y*^ and *Slc26a4*^*C565Y/C565Y*^ mice. Tissue sections were mounted onto the silane-coated glass slides and then were deparaffinized in xylene and rehydrated in ethanol. Tissue were stained with rabbit anti-pendrin, (1:1000; antibody was kindly provided by Dr. Jinsei Jung, Yonsei University College of Medicine, Seoul, Republic of Korea), then incubated with DAPI (1:5000) and Alexa Fluor 488-conjugated goat anti-rabbit IgG (H+L) secondary antibodies (1:200; Thermo Fisher). After incubation, the slides were washed with PBS and mounted with the ProLong Antifade Kit at 25 °C. Images were captured using a laser scanning confocal microscope (LSM880, Zeiss).

### Statistical analysis

Data are shown as means ± SD. Student’s t-test was used for statistical analysis. A p-value of 0.05 was considered to indicate significance.

## Results

### Gross inner ear morphology

Previous studies revealed that pathogenic *Slc26a4* variants could cause enlarged endolymphatic sac, dilated vestibular aqueducts, and inflated scala media [15, 18]. The enlargement of the endolymphatic sac and vestibular aqueducts were observed in *Slc26a4*^*919-2A*>*G/919-2A*>*G*^ mice, but not in *Slc26a4*^+*/*+^ and *Slc26a4*^*C565Y/C565Y*^ mice (Fig. 1C).

### Auditory phenotype evaluation and histology of cochlea

Wild-type mice (i.e., *Slc26a4*^+*/*+^), heterozygous mice (i.e., *Slc26a4*^+*/C565Y*^), and homozygous mice (i.e., *Slc26a4*^*C565Y/C565Y*^) (n = 10 each) were subjected to audiological evaluations at P28 (Fig. 2A). Both *Slc26a4*^+*/C565Y*^ and *Slc26a4*^*C565Y/C565Y*^ mice exhibited normal hearing up to 9 months (data not shown), indicating that the p.C565Y allele is not causal for SNHI in mice.

**Figure 2.**
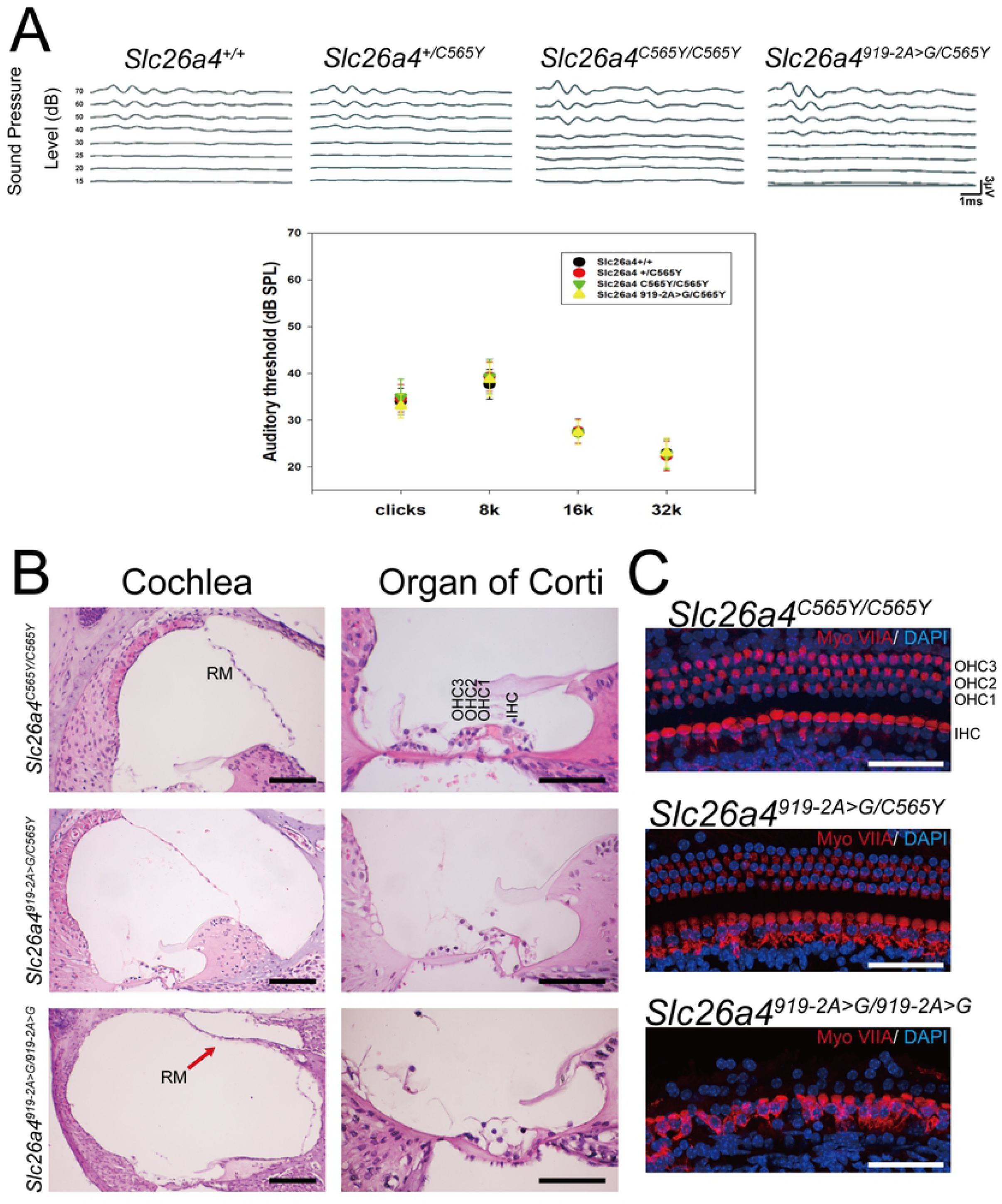
Auditory phenotypes and histology of cochlea. (A) Heterozygous *Slc26a4*^+*/C565Y*^ mice, homozygous *Slc26a4*^*C565Y/C565Y*^ mice, and compound heterozygous *Slc26a4*^*919-2A*>*G/C565Y*^ mice showed normal hearing thresholds across different frequencies, similar to that in wild-type *Slc26a4*^+*/*+^ mice. (B) Histology of the cochlea harvested at 3-month-old. Abnormal histological phenotypes were observed in *Slc26a4*^*919-2A*>*G/919-2A*>*G*^ mice, including dilatation of the scala media (Cochlea panel; RM indicates Reissner’s membrane) and degeneration of the cochlear hair cells (Organ of Corti panel), which were not observed in *Slc26a4*^*C565Y/C565Y*^ and *Slc26a4*^*919-2A*>*G/C565Y*^ mice (bar = 150 μm). (C) Histology of the cochlear hair cells harvested from 3-month-old mice. Myosin VIIA expression was normal in *Slc26a4*^*C565Y/C565Y*^ and *Slc26a4*^*919-2A*>*G/C565Y*^ mice when compared to the diminished expression in *Slc26a4*^*919-2A*>*G/919-2A*>*G*^ mice (bar = 50 μm).

To confirm the pathogenicity of the p.C565Y allele in mice, we further manipulated the *Slc26a4* allele by generating compound heterozygous mice (i.e., *Slc26a4*^*919-2A*>*G/C565Y*^) upon intercrossing the *Slc26a4*^+*/C565Y*^ mice with *Slc26a4*^*919-2A*>*G/919-2A*>*G*^ mice [19]. Similar to the heterozygous mice with c.919-2A>G (i.e., *Slc26a4*^+*/919-2A*>*G*^), *Slc26a4*^*919-2A*>*G/C565Y*^ mice (n = 10) also exhibited normal hearing up to 9 months. This finding indicates that the p.C565Y allele is non-pathogenic and also that the single p.C565Y allele is sufficient to maintain the auditory function in mice.

The histology of the cochlea was also investigated in homozygous mice (i.e., *Slc26a4*^*C565Y/C565Y*^) and compound heterozygous mice (i.e., *Slc26a4*^*919-2A*>*G/C565Y*^). For this, the cochleae of wild-type mice and the profoundly deaf *Slc26a4*^*919-2A*>*G/919-2A*>*G*^ mice were obtained for comparison. Abnormal histological phenotypes that were observed in *Slc26a4*^*919-2A*>*G/919-2A*>*G*^ mice, including dilatation of the scala media (Fig. 2B) and degeneration of the cochlear hair cells (Fig. 2B), were not observed in *Slc26a4*^*C565Y/C565Y*^ and *Slc26a4*^*919-2A*>*G/C565Y*^ mice. Fluorescence confocal microscopy analysis revealed that, unlike in the *Slc26a4*^*919-2A*>*G/919-2A*>*G*^ mice, cochlear hair cells were not degenerated in both *Slc26a4*^*C565Y/C565Y*^ and *Slc26a4*^*919-2A*>*G/C565Y*^ mice (Fig. 2C).

### Vestibular function evaluation and histology of vestibular organs

A total of 60 mice, including 15 *Slc26a4*^+*/*+^, 15 *Slc26a4*^*919-2A*>*G/919-2A*>*G*^, 15 *Slc26a4*^*C565Y/C565Y*^, and 15 *Slc26a4*^*919-2A*>*G/C565Y*^ mice, were subjected to vestibular function evaluation. Similar to the normal audiological phenotypes, homozygous (i.e., *Slc26a4*^*C565Y/C565Y*^) and compound heterozygous (*Slc26a4*^*919-2A*>*G/C565Y*^) mice did not exhibit vestibular deficiencies such as head tilting and circling behavior, and moreover, both these strains performed normally in the swimming and rotarod tests (Fig. 3A). These findings indicate that the single p.C565Y allele is sufficient to maintain normal vestibular function in mice.

**Figure 3.**
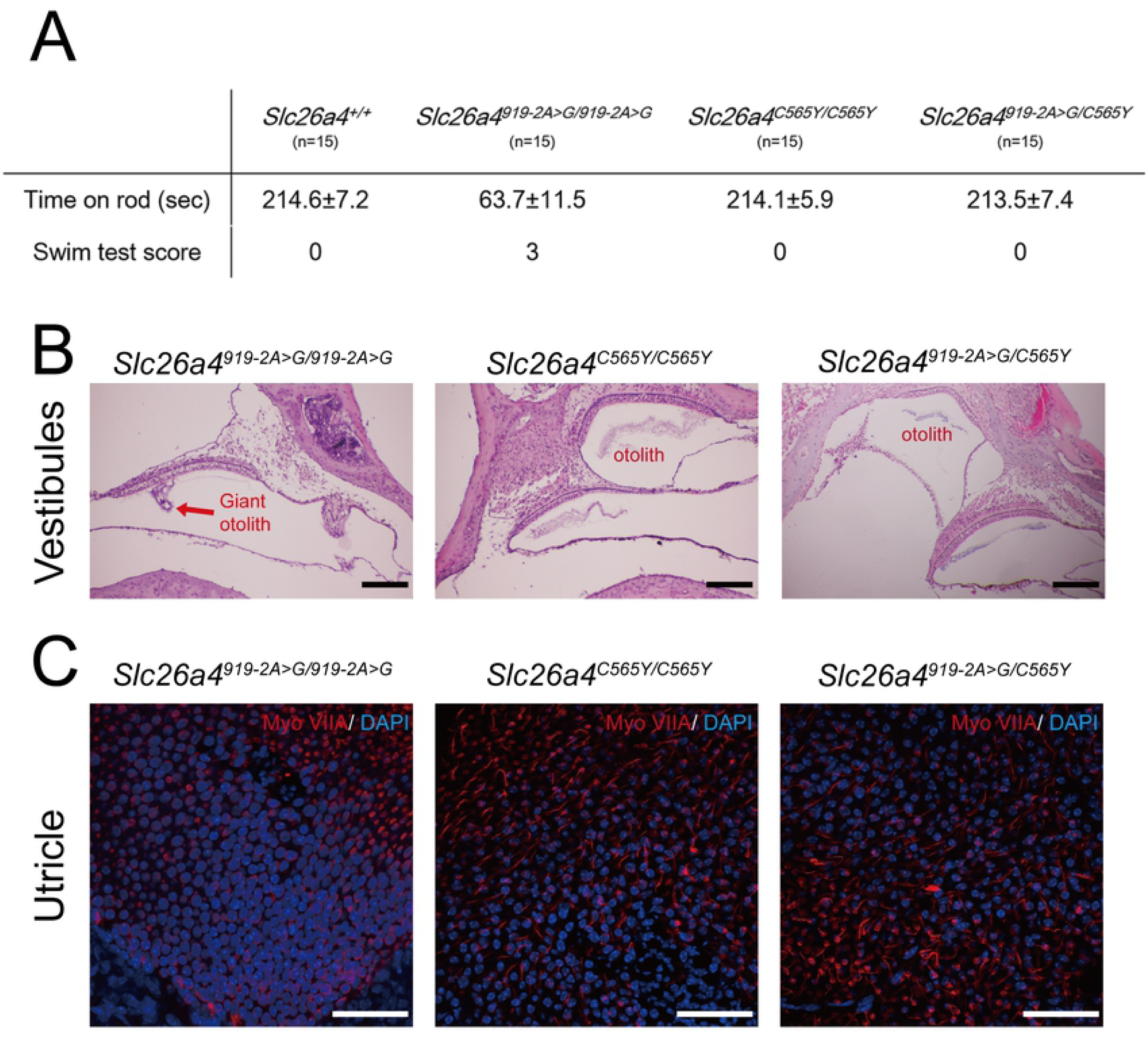
Vestibular phenotypes and histology of vestibular organs. (A) In contrast to the *Slc26a4*^*919-2A*>*G/919-2A*>*G*^ mice, the homozygous *Slc26a4*^*C565Y/C565Y*^ mice, and compound heterozygous *Slc26a4*^*919-2A*>*G/C565Y*^ mice performed well in both, the swimming test and the rotarod test, similar to the wild-type *Slc26a4*^+*/*+^ mice. (B) Histology of the vestibular organs. Giant otoconia was observed in *Slc26a4*^*919-2A*>*G/919-2A*>*G*^ mice, but the otoconia was normal in *Slc26a4*^*C565Y/C565Y*^ and *Slc26a4*^*919-2A*>*G/C565Y*^ mice (bar = 150 μm). (C) Fluorescence confocal microscopy. In contrast to the *Slc26a4*^*919-2A*>*G/919-2A*>*G*^ mice, vestibular hair cells in *Slc26a4*^*C565Y/C565Y*^ mice and *Slc26a4*^*919-2A*>*G/C565Y*^ mice did not degenerate (bar = 50 μm).

Giant otoconia in the vestibular organs (Fig. 3B) and degeneration of the vestibular hair cells (Fig. 3C) were observed in *Slc26a4*^*919-2A*>*G/919-2A*>*G*^ mice, whereas the *Slc26a4*^*C565Y/C565Y*^ and *Slc26a4*^*919-2A*>*G/C565Y*^ mice exhibited normal histology for the vestibular organs (Fig. 3B and 3C).

### Morphology of stria vascularis

Atrophic stria vascularis (SV) with decreased thickness has been documented in mice with various pathogenic *Slc26a4* variants [19-22]. Significant atrophy of SV was observed in *Slc26a4*^*919-2A*>*G/919-2A*>*G*^ mice, but not in *Slc26a4*^*C565Y/C565Y*^ and *Slc26a4*^*919-2A*>*G/C565Y*^ mice (Fig. 4A). There was a significant difference in the SV thickness among the three groups (Fig. 4B, n=3 for each group).

**Figure 4.**
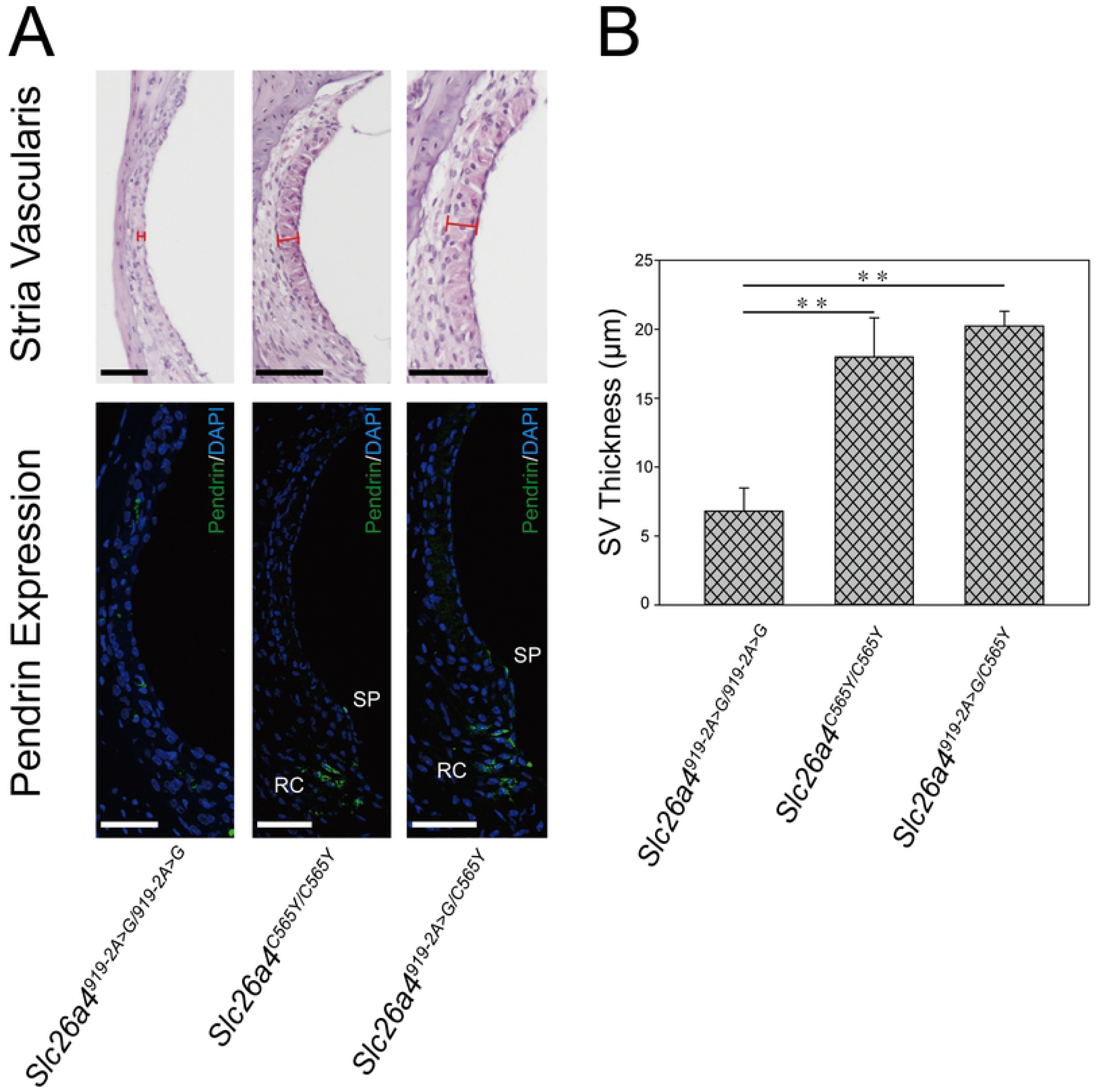
Morphology of stria vascularis and pendrin expression. (A) The histology and the expression of pendrin in the stria vasculasris (SV). Significant atrophy of the SV and poor protein expression of pendrin were observed in *Slc26a4*^*919-2A*>*G/919-2A*>*G*^ mice. By contrast, pendrin was normally distributed in the spiral prominence (SP) and root cells (RC) in the *Slc26a4C*^*565Y/C565Y*^ and *Slc26a4*^*919-2A*>*G/C565Y*^ mice, similar to that in the wild-type mice. Tissues were harvested from 3-month-old mice (bar = 50μm). (B) Quantification data of the SV thickness. The SV thickness in each group of mice was calculated (n=3). In *Slc26a4*^*C565Y/C565Y*^ and *Slc26a4*^*919-2A*>*G/C565Y*^ mice, the SV was thicker than that in *Slc26a4*^*919-2A*>*G/919-2A*>*G*^ mice, in which the SV was atrophic (\*\**p* <0.01).

We then examined the expression of pendrin in the cochlea of *Slc26a4*^*C565Y/C565Y*^ mice and *Slc26a4*^*919-2A*>*G/C565Y*^ mice by immunolocalization assay (Fig. 4A). In both strains, pendrin was normally distributed in root cells and apical membranes of spiral prominence surface epithelial cells, as in wild-type mice [23], indicating that the expression of pendrin was unaffected in mice with the p.C565Y variant.

## Discussion

In this study, we generated a transgenic mouse model harboring the p.C565Y variant of the *Slc26a4* gene to examine the pathogenicity of the p.C565Y variant in mice. Phenotypic characterization revealed that both homozygous (i.e., *Slc26a4*^*919-2A*>*G/C565Y*^) and compound heterozygous (i.e., *Slc26a4*^*919-2A*>*G/C565Y*^) mice exhibit normal hearing and balance, which was further confirmed by the normal inner ear morphology in histological studies. These findings indicate that the p.C565Y variant of *Slc26a4* gene is non-pathogenic for mice.

*SLC26A4* p.C565Y variant has been documented as a pathogenic variant in several previous reports [9, 24-28]. Using the Varsome platform [29], the *SLC26A4* p.C565Y variant meets the ACMG criteria for PM1, PM2, PP2, PP3, and PP5. It is located at a functional domain of the *SLC26A4* gene rich in pathogenic mutational hotspots (PM1), and exhibits minor allelic frequency (MAF) <0.001 across various populations in the gnomAD browser (PM2). The *SLC26A4* gene also possess large proportion of pathogenic missense variants that are recorded in Uniport (PP2). In some databases (ClinVar, DVD, and Uniport) and in several prediction algorithms (SIFT, DANN, EIGEN, FATHMM-MKL, MutationTaster, etc.), the p.C565Y variant is annotated to be a pathogenic variant (PP3 and PP5), and the SIFT score suggests that it is damaging (0.004), while the polyphen2 score suggests that it is moderately pathogenic (0.714). Therefore, the *SLC26A4* p.C565Y variant can be classified as “likely pathogenic” according to the ACMG guidelines, indicating that its pathogenicity in humans might differ from that in mice.

The mutation landscape of *SLC26A4* gene differs significantly among different ethnic groups [25, 28, 30, 31]. Previous clinical studies have revealed inconsistent results in regard to the correlation between *SLC26A4* genotypes and phenotypes [27, 32, 33]. In several studies, the authors did not identify any phenotypic differences among patients with different *SLC26A4* variants [26, 34]. However, it has been recently reported that some variants might lead to progression of hearing loss [35, 36], and some *SLC26A4* genotypes might be associated with a normal thyroid phenotype and less severe hearing loss [37].

Consistent with these recent clinical reports, studies based on cell-line experiments demonstrated that different *SLC26A4* variants have different pathogenic mechanisms. Some variants, such as p.H723R, p.L236P, and p.T721M, are shown to be associated with defective protein expression or trafficking, whereas some variants, such as p.K369E, and p.S166N, are shown to be associated with normal protein expression but impaired protein function [7-9]. Notably, even among each subgroup, there seems to be a gradient of pathogenicity. For instance, the trafficking of pendrin in p.H723R variant could be rescued by salicylate and temperature manipulation, whereas pendrin in p.L236P variant could not be rescued. The p.C565Y variant seems to be non-pathogenic in cell lines as pendrin with p.C565Y is expressed correctly in the plasma membranes in both HEK 293 [7, 8] and COS-7 cell lines [9], and is also shown to exhibit normal protein function in Xenopus oocytes [9].

Similarly, different *SLC26A4* variants are also shown to be associated with different phenotypic severities in animal models. To date, several mouse models have been generated, including the knock-out *Slc26a4*^*-/-*^ mice [10], *Slc26a4*^*loop/loop*^ mice with the p.S408F variant [11], *Slc26a4*^*919-2A*>*G9/919-2A*>*G*^ mice with the c.919-2 A>G variant [19], *Slc26a4*^*H723R/H723R*^ mice with the p.H723R variant [13], *Slc26a4*^*L236P/L236P*^ mice with the p.L236P variant [14], and the conditional knock-outs Tg[E]; Tg[R]; *Slc26a4*^*Δ/Δ*^ mice [15] (Table 1). The *Slc26a4*^*-/-*^, *Slc26a4*^*loop/loop*^, and *Slc26a4*^*919-2A*>*G9/919-2A*>*G*^ mice exhibit congenital profound SNHI, which represents the most severe symptom in the phenotypic spectrum, whereas the *Slc26a4*^*H723R/H723R*^ mice exhibit normal hearing, representing the least severe symptom in the phenotypic spectrum. The *Slc26a4*^*L236P/L236P*^ and the Tg[E], Tg[R], and *Slc26a4*^*Δ/Δ*^ mice revealed an intermediate phenotype, such that some residual hearing could be recorded in both strains.

**Table 1.**
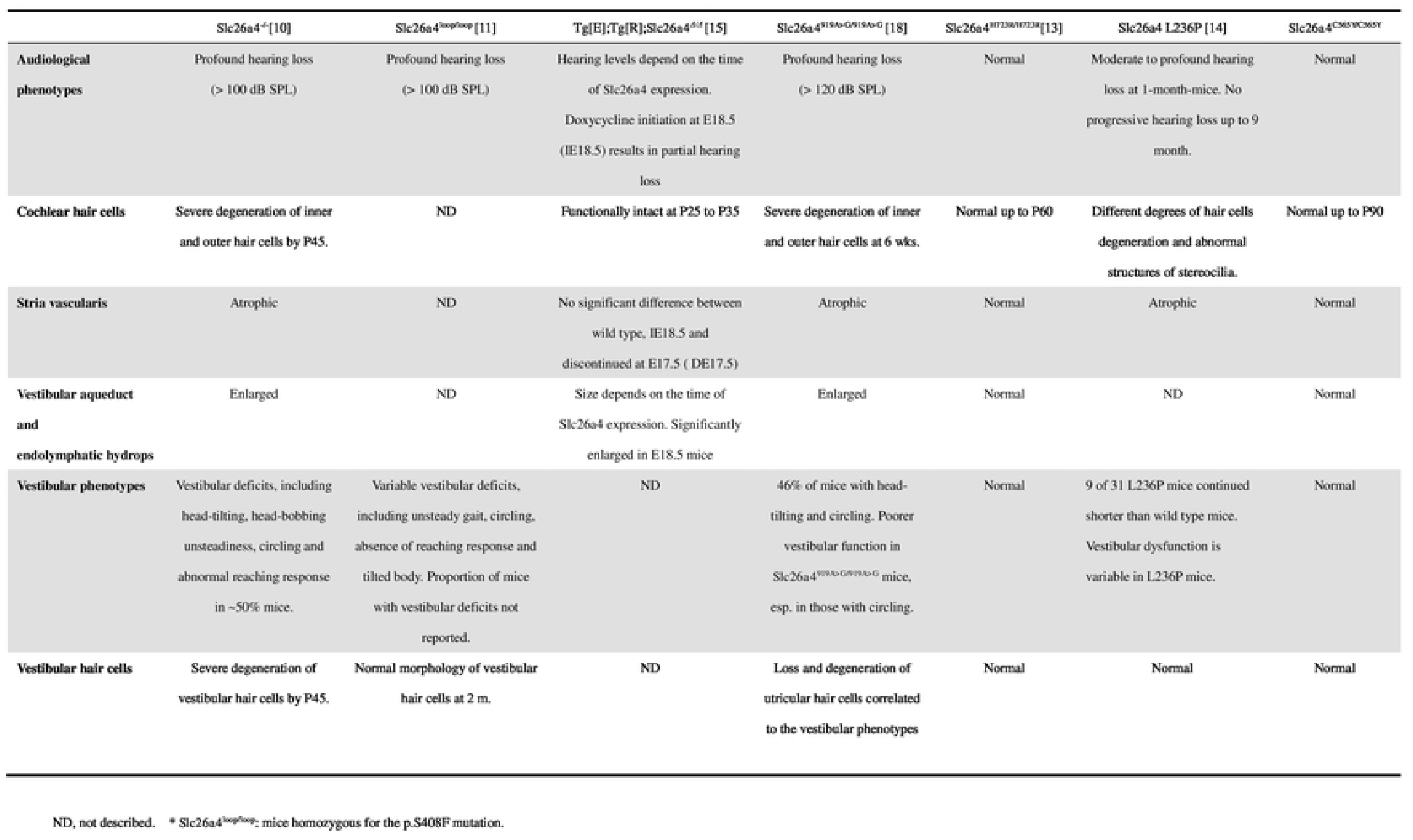
Comparison of phenotypes among mouse strains with different *Slc26a4* variants.

Interestingly, there seems to be some correlation between the pathogenicity in the cell lines and animals. Variants with stronger pathogenicity in cell lines, such as p.L236P, correlates with the presence of phenotypes in transgenic mice, whereas variants with weaker or no pathogenicity in cell lines, such p.H723R and p.C565Y, correlates with the absence of phenotypes in transgenic mice. This correlation might be helpful for researchers in selecting *SLC26A4* variants as potential subjects for generating the transgenic mouse models accordingly.

Despite the presence of these correlations in the pathogenicity of *SLC26A4* variants, it should be noted that the cell-line and transgenic mice experiments still possess some limitations that preclude them from being satisfactory experimental models for *SLC26A4*-related SNHI. First of all, some variants that have been supported by strong evidence to be clinically pathogenic, such as p.H723R or p.C565Y, did not reflect any abnormality in protein expression or function in cell lines [8], or audiovestibular phenotypes in transgenic mice [13]. It has also been reported that the trafficking/glycosylation process might significantly differ between mice and humans [38], which might explain the absence of phenotypes in mice with p.C565Y and p.H723R variants of *Slc26a4* gene. Second, even for transgenic mice with positive phenotypes, there are still valid discrepancies in the audiovestibular features that are reported between mice and humans. For instance, the congenital profound SNHI reported in *Slc26a4*^*-/-*^, *Slc26a4*^*loop/loop*^, and *Slc26a4*^*919-2A*>*G9/919-2A*>*G*^ mice is too severe as compared to their phenotype in human counterpart. Meanwhile, although the *Slc26a4*^*L236P/L236P*^ and Tg[E]; Tg[R]; *Slc26a4*^*Δ/Δ*^ mice showed some residual hearing, neither could perfectly recapitulate the fluctuating or progressive disease nature in humans. Further studies are required to refine the cell-line or transgenic mouse models to investigate and to better understand the *SLC26A4*-related SNHI.

In conclusion, using a genotype-driven approach, we generated a knock-in mouse model that segregated the deafness-associated *SLC26A4* p.C565Y variant in humans. Subsequent phenotypic characterization revealed that mice with the *Slc26a4* p.C565Y variant exhibit normal audiovestibular phenotypes and inner ear morphology, indicating that the pathogenicity associated with specific *SLC26A4* variants might differ between humans and mice. Therefore, with regard to the research of *SLC26A4*-related SNHI, caution should be taken when extrapolating the results of animal studies to humans.

## Acknowledgements

We thank the staff of the imaging core at the First Core Lab, National Taiwan University College of Medicine, for technical assistance. We also thank National Taiwan University Hospital (NTUH.108-T14) and Taipei Veterans General Hospital - National Taiwan University Hospital Joint Research Program (VN109-15 & VN108-09) for the grant support.

## References

[1] Wolf, A. et al. A novel mutation in SLC26A4 causes nonsyndromic autosomal recessive hearing impairment. Otology & Neurotology 38, 173–179 (2017).

[2] Han, M. et al. A quantitative cSMART assay for noninvasive prenatal screening of autosomal recessive nonsyndromic hearing loss caused by GJB2 and SLC26A4 mutations. Genetics in Medicine 19, 1309 (2017).

[3] Everett, L. A., Morsli, H., Wu, D. K. & Green, E. D. Expression pattern of the mouse ortholog of the Pendred’s syndrome gene (Pds) suggests a key role for pendrin in the inner ear. Proceedings of the National Academy of Sciences 96, 9727–9732 (1999).

[4] Pelzl, L. et al. DOCA sensitive pendrin expression in kidney, heart, lung and thyroid tissues. Cellular Physiology and Biochemistry 30, 1491–1501 (2012).

[5] Li, X. C. et al. A mutation in PDS causes non-syndromic recessive deafness. Nature genetics 18, 215 (1998).

[6] Mori, T., Westerberg, B. D., Atashband, S. & Kozak, F. K. Natural history of hearing loss in children with enlarged vestibular aqueduct syndrome. Journal of Otolaryngology--Head & Neck Surgery 37 (2008).

[7] Yoon, J. S. et al. Heterogeneity in the processing defect of SLC26A4 mutants. J Med Genet 45, 411–419, doi:10.1136/jmg.2007.054635 (2008).

[8] Ishihara, K. et al. Salicylate restores transport function and anion exchanger activity of missense pendrin mutations. Hear Res 270, 110–118, doi:10.1016/j.heares.2010.08.015 (2010).

[9] Choi, B. Y. et al. Hypo-functional SLC26A4 variants associated with nonsyndromic hearing loss and enlargement of the vestibular aqueduct: genotype-phenotype correlation or coincidental polymorphisms? Hum Mutat 30, 599–608, doi:10.1002/humu.20884 (2009).

[10] Everett, L. A. et al. Targeted disruption of mouse Pds provides insight about the inner-ear defects encountered in Pendred syndrome. Human molecular genetics 10, 153–161 (2001).

[11] Dror, A. A. et al. Calcium oxalate stone formation in the inner ear as a result of an Slc26a4 mutation. Journal of Biological Chemistry 285, 21724–21735 (2010).

[12] Lu, Y. C. et al. Establishment of a knock-in mouse model with the SLC26A4 c.919-2A>G mutation and characterization of its pathology. PLoS One 6, e22150, doi:10.1371/journal.pone.0022150 (2011).

[13] Lu, Y.-C. et al. Differences in the pathogenicity of the p. H723R mutation of the common deafness-associated SLC26A4 gene in humans and mice. PloS one 8, e64906 (2013).

[14] Wen, Z. et al. A knock-in mouse model of Pendred syndrome with Slc26a4 L236P mutation. Biochem Biophys Res Commun 515, 359–365, doi:10.1016/j.bbrc.2019.05.157 (2019).

[15] Choi, B. Y. et al. Mouse model of enlarged vestibular aqueducts defines temporal requirement of Slc26a4 expression for hearing acquisition. The Journal of clinical investigation 121 (2011).

[16] Hardisty-Hughes, R. E., Parker, A. & Brown, S. D. A hearing and vestibular phenotyping pipeline to identify mouse mutants with hearing impairment. Nature protocols 5, 177 (2010).

[17] Isgrig, K. et al. Gene therapy restores balance and auditory functions in a mouse model of Usher syndrome. Molecular Therapy 25, 780–791 (2017).

[18] Takeda, H. et al. prenatal electroporation-mediated gene transfer restores Slc26a4 knock-out mouse hearing and vestibular function. Scientific Reports 9, 1–12 (2019).

[19] Lu, Y.-C. et al. Establishment of a knock-in mouse model with the SLC26A4 c. 919-2A> G mutation and characterization of its pathology. PLoS One 6, e22150 (2011).

[20] Kim, H.-M. & Wangemann, P. Epithelial cell stretching and luminal acidification lead to a retarded development of stria vascularis and deafness in mice lacking pendrin. PloS one 6 (2011).

[21] Ito, T. et al. Slc26a4-insufficiency causes fluctuating hearing loss and stria vascularis dysfunction. Neurobiology of disease 66, 53–65 (2014).

[22] Royaux, I. E. et al. Localization and functional studies of pendrin in the mouse inner ear provide insight about the etiology of deafness in pendred syndrome. Journal of the Association for Research in Otolaryngology 4, 394–404 (2003).

[23] Wangemann, P. et al. Loss of KCNJ10 protein expression abolishes endocochlear potential and causes deafness in Pendred syndrome mouse model. BMC medicine 2, 30 (2004).

[24] Pryor, S. P. et al. SLC26A4/PDS genotype-phenotype correlation in hearing loss with enlargement of the vestibular aqueduct (EVA): evidence that Pendred syndrome and non-syndromic EVA are distinct clinical and genetic entities. J Med Genet 42, 159–165, doi:10.1136/jmg.2004.024208 (2005).

[25] Tsukamoto, K. et al. Distribution and frequencies of PDS (SLC26A4) mutations in Pendred syndrome and nonsyndromic hearing loss associated with enlarged vestibular aqueduct: a unique spectrum of mutations in Japanese. European journal of human genetics 11, 916 (2003).

[26] Miyagawa, M., Nishio, S.-y. & Usami, S.-i. Mutation spectrum and genotype–phenotype correlation of hearing loss patients caused by SLC26A4 mutations in the Japanese: a large cohort study. Journal of human genetics 59, 262 (2014).

[27] Suzuki, H. et al. Clinical characteristics and genotype–phenotype correlation of hearing loss patients with SLC26A4 mutations. Acta otolaryngologica 127, 1292–1297 (2007).

[28] Campbell, C. et al. Pendred syndrome, DFNB4, and PDS/SLC26A4 identification of eight novel mutations and possible genotype–phenotype correlations. Human mutation 17, 403–411 (2001).

[29] Kopanos, C. et al. VarSome: the human genomic variant search engine. Bioinformatics 35, 1978–1980, doi:10.1093/bioinformatics/bty897 (2019).

[30] Song, M.-J. et al. Estimation of carrier frequencies of six autosomal-recessive Mendelian disorders in the Korean population. Journal of human genetics 57, 139 (2012).

[31] Tsukada, K., Nishio, S.-y., Hattori, M. & Usami, S.-i. Ethnic-specific spectrum of GJB2 and SLC26A4 mutations: their origin and a literature review. Annals of Otology, Rhinology & Laryngology 124, 61S–76S (2015).

[32] Azaiez, H. et al. Genotype–phenotype correlations for SLC26A4-related deafness. Human genetics 122, 451–457 (2007).

[33] Wu, C.-C., Chen, P.-J. & Hsu, C.-J. Specificity of SLC26A4 mutations in the pathogenesis of inner ear malformations. Audiology and Neurotology 10, 234–242 (2005).

[34] Wu, C.-C. et al. Phenotypic analyses and mutation screening of the SLC26A4 and FOXI1 genes in 101 Taiwanese families with bilateral nonsyndromic enlarged vestibular aqueduct (DFNB4) or Pendred syndrome. Audiology and Neurotology 15, 57–66 (2010).

[35] Lee, H. et al. Correlation between genotype and phenotype in patients with bi-allelic SLC26A4 mutations. Clinical genetics 86, 270–275 (2014).

[36] Rah, Y. C. et al. Audiologic presentation of enlargement of the vestibular aqueduct according to the SLC 26 A 4 genotypes. The Laryngoscope 125, E216–E222 (2015).

[37] Chao, J. R. et al. SLC26A4-linked CEVA haplotype correlates with phenotype in patients with enlargement of the vestibular aqueduct. BMC Med Genet 20, 118, doi:10.1186/s12881-019-0853-4 (2019).

[38] Zhang, B.-Y. L. a. J. Null mutations in human and mouse orthologs frequently result in different phenotypes. Proceeding of the National Academy of Sciences of the United States of America 105, 6 (2008).

